# The concreteness effect from memory illusions’ perspective: the DIM-HA effect

**DOI:** 10.1101/2023.07.06.548014

**Authors:** Alejandro Marín-Gutiérrez, Emiliano Díez Villoria, Ana María González

**Affiliations:** Facultad de Educación y Psicología, Universidad del Atlántico Medio, Gran Canaria, Spain.; Departamento de Psicología Básica, Psicobiología y Metodología de las Ciencias del Comportamiento. Universidad de Salamanca, Salamanca, Spain; Instituto Universitario de Integración en la Comunidad (INICO), Universidad de Salamanca, Salamanca, Spain

**Author notes:** These authors contributed equally to this work. Corresponding Author: Alejandro Marín-Gutiérrez.

**Keywords:** DRM paradigm, Concreteness Effect, QDR theory, Associative Activation Theory, Dim-Ha Effect.

## Abstract

A vast body of evidence has shown that concrete nouns are processed faster and more accurately than abstract nouns in a variety of cognitive tasks. This phenomenon is widely known as the concreteness effect and explanations for its occurrence seem to reflect differences in processing and organization between both type of concepts. While there is considerable evidence to support this concrete effect, the nature of these differences is still controversial. In developing this explanation, we have proposed a relatively different approach from a false memory perspective using the DRM paradigm. Three different experiments were created to address the importance of association in creating concrete and abstract false memories. Results showed that false recognition rates differed significantly between concrete and abstract critical words when they were associated strongly with their respective lists, which led to a higher proportion of abstract false alarms both in behavioral and electrophysiological experiments. The principal outcomes were discussed in terms of theories of associative activation and qualitatively different representation.

## Introduction

Concepts that allude to physical objects (such as puppy) are considered concrete, whereas concepts that refer to non-physical entities (such as idea) are considered abstract. Between these two categories, there is a continuum whereby all conceptual information is placed according to the links that the concepts share with the physical entities. In this context, the *concreteness effect* is defined as the processing advantage, in terms of speed and accuracy, for concrete over abstract concepts. Such concrete over abstract processing advantages has been demonstrated repeatedly from behavioral tasks (Bleasdale, 1987; Crutch at al. 2009; Duñabeitia et al., 2009; Schwanenflugel et al,1992), neuroimaging techniques (van Schie et al., 2005; Wang et al. 2010), and neuropsychological patients (Breedin et al. Coslett, 1994; Crutch et al., 2009; Crutch & Warrington, 2005, 2006). Despite robust observation, no convincing explanation exists for its functional properties or structural characteristics.

Paivio and Csapo proposed one of the first explanations for the concreteness effect in their *Dual Coding Theory* (Paivio & Csapo, 1969). The authors hypothesize that the advantage of processing concrete over abstract terms is that concrete concepts are coded by both a verbal and an image system, whereas abstract concepts are coded only by the verbal system. Thus, concrete words are processed via two pathways, providing them with a higher imageability (Boles, 1983; Richardson, 1976).

Inspired by this dual coding logic, Schwanenflugel et al., (1992) discussed three different hypotheses that might explain the concreteness effect from a psycholinguistic point of view. The first one, known as the *Automatic-Imagery hypothesis*, states that concrete concepts, unlike abstract ones, are related to sensory experiences, which automatically makes them easier to remember as they are represented in two ways: verbally and imaginably. The second one is the *Strategic-Imagery hypothesis*, which claims that imagery is used deliberately when it is considered a helpful strategy. The final hypothesis is the *Context-Availability hypothesis,* which proposes that prior knowledge sets a frame in the processing of concrete and abstract concepts. Concrete concepts are supposed to be easier comprehended and recalled because related contextual knowledge is more accessible (e.g., the words have a higher context-availability). The results of the study of Schwanenflugel et al. (1992), seem to support more the *Strategic-Imagery* hypothesis.

A different finding came from Kousta et al., (2011). The authors found that when controlling for imageability and context availability, there is a processing advantage for abstract but not for concrete words. Further experiments showed that this abstractness effect must have been produced due to the affective associations of the words, as abstract words tend to have higher emotional valence scores than concrete words.

To add more evidence for explaining the concreteness effect, Crutch & Warrington (2005) investigated the comprehension of abstract and concrete words from a neuropsychological approach. They conducted different word-matching tasks with a globally aphasic patient. Based on the results, they proposed the existence of qualitative differences in the representation of abstract and concrete concepts. According to them, concrete words are primarily organized based on taxonomic similarity, whereas abstract concepts seem to rely on the associative connections among them. As concreteness is considered a continuum, a concept can have both associative and categorical connections in different proportions. An associative network might provide the flexibility that is necessary to involve all the different, context-dependent meanings that abstract concepts need to be successfully processed. Thus, this stablished the birth of the *Qualitatively Different Representational* theory (QDR).

Further evidence for this QDR framework was found in a study with healthy participants using the visual-world paradigm (Duñabeitia et al. 2009). After hearing an abstract word, the participants’ attention tended to be captured more and earlier by a picture of a semantically associated object (among distractors) than after hearing a concrete word. Their results were interpreted as supporting evidence for the different organizational principles described by the QDR framework. Similarly, Crutch et al. (2009) also tested healthy participants who performed various odd-one-out tasks. They tended to process the connections between concrete words faster when they were similar than when they were associated. The reverse pattern was shown with abstract words.

Additionally, Mkrtychian et al. (2021) reported differences in behavioral and neurophysiological processing between concrete and abstract words that appear immediately after controlled acquisition, confirming that distinct neurocognitive mechanisms underlie these semantic representations.

The analysis of the reviewed literature seems to highlight the importance of the association strength, especially for processing abstract concepts. To further explore this assumption, the research we propose is aimed at contributing to this debate by using the Deese-Roediger-McDermott (DRM) paradigm (Deese, 1959; Roediger & McDermott, 1995), this is, from the false memory perspective. Although there has been plenty of evidence of the concreteness effect from the memory perspective, the analysis of memory disruption in healthy people should represent a convenient field of study to provide convergent evidence in this matter.

### The DRM paradigm

The DRM paradigm is based on the presentation of word lists to participants, each of which contains a keyword (or critical lure) that is never presented during the study phase. Later, during the test phase, the researcher presents the lure, thereby activating the false memory effect. In other words, when participants are later asked to recall the presented items, they frequently recall the critical words incorrectly. In the following paragraphs, various hypotheses regarding the causes of these false memories in the DRM paradigm will be discussed.

### The Associative-activation theory

In Gallo’s review (2006; 2010), the associative-activation theory describes that when participants study the list of words (e.g., *puppy, bark, pet, cat, friend*), they activate not only the word’s representation but also that activation spreads throughout associated concepts. At the test, the pre-activated representations (such as the critical word *“dog”*) are more available and more prone to be selected as part of the studied material (e.g., producing a false memory). This theory was part of the descriptions of the Activation/Monitoring Framework (e.g., Roediger et al. 2001).

This theory relies on the concept of spreading activation: when a word is studied, its representation within the conceptual system is activated, and this activation spreads out towards the representation of associated concepts that are placed relatively close in the semantic network. To produce memory illusions, this theory proposes that participants may mistakenly believe that a non-presented associate occurred in the list because the lure items have more activation, being this a criterion of selection. In other words, the more activation shows a concept, the more familiar it sounds.

### Gist theory

Fuzzy Trace Theory is a psychological framework that explains how memories are formed and retrieved. Although it does not focus specifically on false memories, it can provide insight into how they can occur. According to Fuzzy Trace Theory, when we encode and store information, we produce both verbatim traces (exact details) and gist traces (meaning or essence). During memory retrieval, reliance on essence traces rather than verbatim traces can lead to the formation of false memories.

*Fuzzy Trace Theory* explains false memories as follows:

*Gist-Based Encoding:* during memory encoding, people tend to derive the gist or meaning of an event rather than meticulously storing every detail. Specific details may be forgotten or altered because of this reliance on gist processing, resulting in memory distortions. False memories can occur when the essence of an event is remembered correctly, but the specific details are incorrectly reconstructed or influenced by external information.

When recalling memories, people frequently rely on essence traces rather than exact verbatim traces. This process entails the reconstruction of memories using general knowledge and schema. Individuals with false memories may unconsciously fill in missing details with plausible or suggested information, leading them to believe that the reconstructed memory is authentic.

*Fuzzy Trace Theory* also highlights the importance of suggestibility and misinformation in the formation of false memories. The reconstruction process can be influenced by external information, such as leading inquiries or misleading suggestions. If erroneous information corresponds to the essence or general significance of an event, it may be stored in memory, resulting in a false recall.

The reliance on paraphrase processing during memory encoding and retrieval, according to *Fuzzy Trace Theory*, may result in the formation of false memories. The reconstruction of memories based on general meaning, suggestibility, and misinformation can contribute to the formation of deceptive memories. False memories are complex phenomena influenced by a variety of cognitive and social factors, and *Fuzzy Trace Theory* provides a framework for comprehending some of these processes.

### False memories and concreteness

In the case of the relationship between concreteness and false memories, the question that arises is whether there are differences between false memories of concrete and abstract words. There is not much information regarding this topic, except the study of Pérez-Mata et al. (2002), who used the DRM paradigm to compare false memory rates of concrete and abstract words. They found that concrete items were recalled correctly more often than abstract items and that false recall of non-presented words was higher for abstract lists than for concrete lists.

Another study where concreteness was addressed, although not directly, was Roediger et al.’s (2001), but the authors didn’t obtain a significant effect of concreteness on false memory. In this study, the authors mentioned that the most relevant factor in determining the production of false memory was association strength. This result is interesting because if we delve deeper into the concreteness effect theories, they affirm that association has a special and differential role in the processing of concrete vs. abstract concepts. Specifically, they mentioned the *Backward Association Strength* (BAS) of the lists which had already been reported by Deese (1959). BAS describes the “average tendency for words in the study list to elicit the critical item on a free association test” (Roediger et al., 2001). Therefore, lists with high BAS correlate with higher rates of false recall (Deese, 1959; Roediger, et al., 2001), as they probably promote activation of the critical concept more than those lists with low BAS.

### The current study

In the context of the literature reviewed, the main goal of the present study was to explore the differences in false recognition of critical abstract words and critical concrete words, while determining the possible differential role of BAS. We hypothesized that false recognition rates should be higher for abstract critical words with high backward association strength.

To have a broad approach to this phenomenon, we included both behavioral and electrophysiological measures.

## Experiment 1

As mentioned in the introduction, the role of concreteness in false memory production is still a matter of controversy. Besides, the effect has been explored in recall, but not in recognition memory, which relies on different cognitive processes (Gillund & Shiffrin, 1984). To explore the relationship between association strength and concreteness over false recognition, we created world lists weakly or strongly associated both with a concrete or abstract critical lure. As it was established in the introduction section, we hypothesized that a concreteness effect would emerge, and association strength would modulate this effect differently for concrete and abstract lists.

## Method

### Participants

24 psychology students (23 females and one male) of the University of Salamanca took part in the experiment in exchange for course credits. Their mean age was 20,54 years. In all the experiments of this research, participants were asked to sign a written consent form. Also, all the participants were recruited from the curse 2014/2015.

### Design and materials

A 2 × 2 within-participants design was used with concreteness (abstract vs. concrete) and associative strength (high BAS vs. low BAS) as independent variables. The proportion of correct and false recognition were the dependent variables.

The word lists were created around 32 critical lure words (16 abstract and 16 concrete; mean: 2.69 and 5.92; range: 1.38-3.38 and 5.03-6.78 respectively) that were taken from the Algarabel study (1996). Additionally, indexes of word frequency, orthographic neighborhood, familiarity, word length, and imageability were obtained for the 32 critical lures. Subsequently, the 32 critical words were introduced into NIPE, a web tool from the University of Salamanca’s research team on memory and cognition where up to date it is possible to obtain association norms for 4.051 words (Díez et al.2006). For every critical lure, we obtained twelve backward associates to build the experimental and distractor material. Table 1 contains the psycholinguistic properties for each critical lure, as well as the association strength average for each list.

**Table 1.**
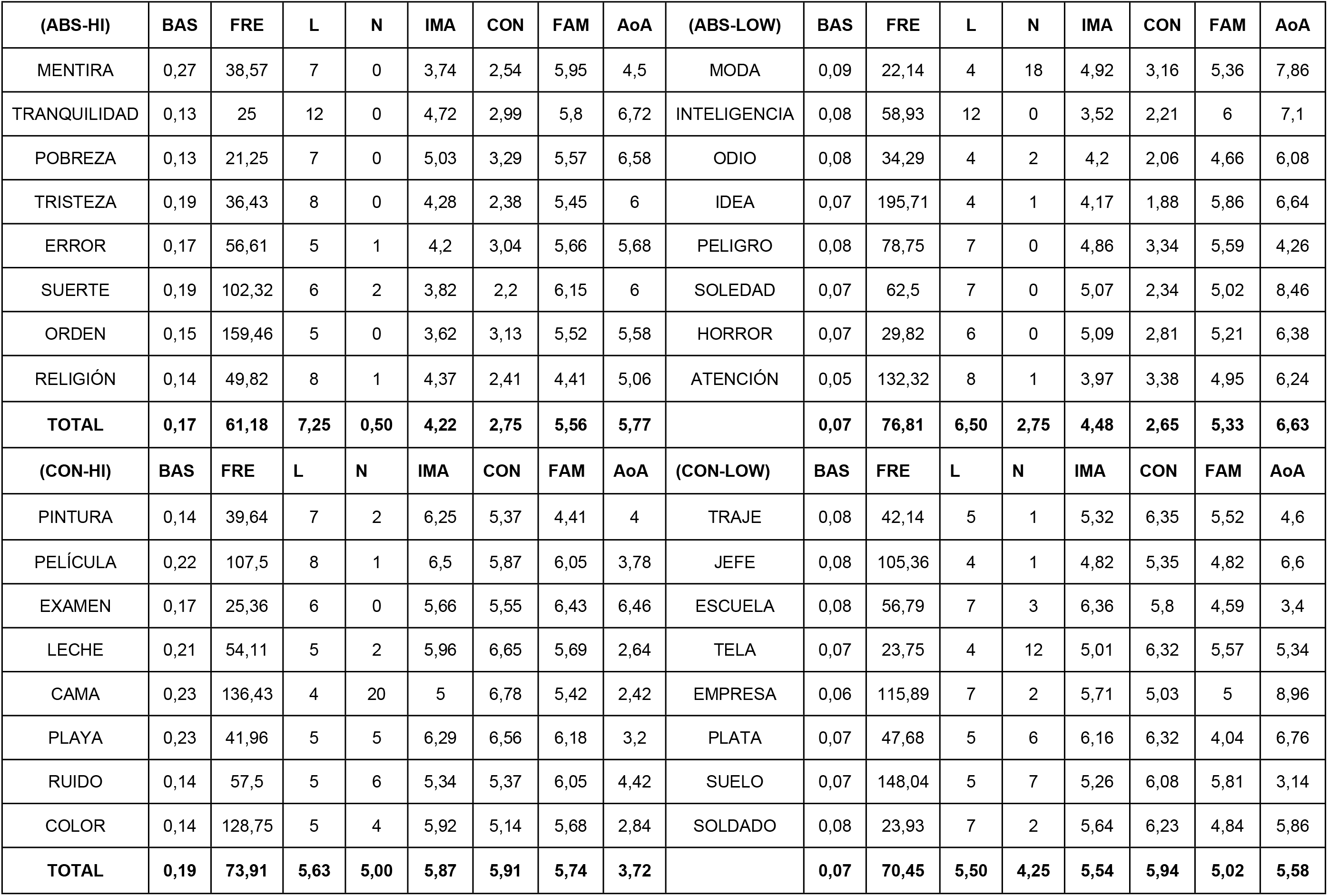
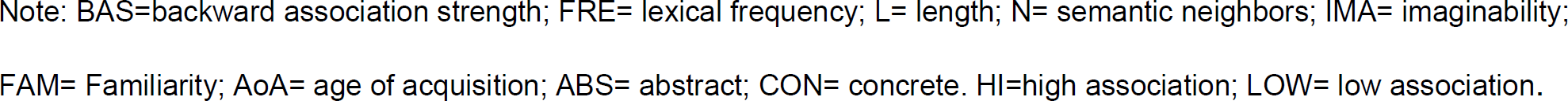
Psycholinguistic properties of the list and critical lures.

| (ABS-HI) | BAS | FRE | L | N | IMA | CON | FAM | AoA | (ABS-LOW) | BAS | FRE | L | N | IMA | CON | FAM | AoA |
| --- | --- | --- | --- | --- | --- | --- | --- | --- | --- | --- | --- | --- | --- | --- | --- | --- | --- |
| MENTIRA | 0,27 | 38,57 | 7 | 0 | 3,74 | 2,54 | 5,95 | 4,5 | MODA | 0,09 | 22,14 | 4 | 18 | 4,92 | 3,16 | 5,36 | 7,86 |
| TRANQUILIDAD | 0,13 | 25 | 12 | 0 | 4,72 | 2,99 | 5,8 | 6,72 | INTELIGENCIA | 0,08 | 58,93 | 12 | 0 | 3,52 | 2,21 | 6 | 7,1 |
| POBREZA | 0,13 | 21,25 | 7 | 0 | 5,03 | 3,29 | 5,57 | 6,58 | ODIO | 0,08 | 34,29 | 4 | 2 | 4,2 | 2,06 | 4,66 | 6,08 |
| TRISTEZA | 0,19 | 36,43 | 8 | 0 | 4,28 | 2,38 | 5,45 | 6 | IDEA | 0,07 | 195,71 | 4 | 1 | 4,17 | 1,88 | 5,86 | 6,64 |
| ERROR | 0,17 | 56,61 | 5 | 1 | 4,2 | 3,04 | 5,66 | 5,68 | PELIGRO | 0,08 | 78,75 | 7 | 0 | 4,86 | 3,34 | 5,59 | 4,26 |
| SUERTE | 0,19 | 102,32 | 6 | 2 | 3,82 | 2,2 | 6,15 | 6 | SOLEDAD | 0,07 | 62,5 | 7 | 0 | 5,07 | 2,34 | 5,02 | 8,46 |
| ORDEN | 0,15 | 159,46 | 5 | 0 | 3,62 | 3,13 | 5,52 | 5,58 | HORROR | 0,07 | 29,82 | 6 | 0 | 5,09 | 2,81 | 5,21 | 6,38 |
| RELIGIÓN | 0,14 | 49,82 | 8 | 1 | 4,37 | 2,41 | 4,41 | 5,06 | ATENCIÓN | 0,05 | 132,32 | 8 | 1 | 3,97 | 3,38 | 4,95 | 6,24 |
| **TOTAL** | **0,17** | **61,18** | **7,25** | **0,50** | **4,22** | **2,75** | **5,56** | **5,77** |  | **0,07** | **76,81** | **6,50** | **2,75** | **4,48** | **2,65** | **5,33** | **6,63** |
| (CON-HI) | BAS | FRE | L | N | IMA | CON | FAM | AoA | (CON-LOW) | BAS | FRE | L | N | IMA | CON | FAM | AoA |
| PINTURA | 0,14 | 39,64 | 7 | 2 | 6,25 | 5,37 | 4,41 | 4 | TRAJE | 0,08 | 42,14 | 5 | 1 | 5,32 | 6,35 | 5,52 | 4,6 |
| PELÍCULA | 0,22 | 107,5 | 8 | 1 | 6,5 | 5,87 | 6,05 | 3,78 | JEFE | 0,08 | 105,36 | 4 | 1 | 4,82 | 5,35 | 4,82 | 6,6 |
| EXAMEN | 0,17 | 25,36 | 6 | 0 | 5,66 | 5,55 | 6,43 | 6,46 | ESCUELA | 0,08 | 56,79 | 7 | 3 | 6,36 | 5,8 | 4,59 | 3,4 |
| LECHE | 0,21 | 54,11 | 5 | 2 | 5,96 | 6,65 | 5,69 | 2,64 | TELA | 0,07 | 23,75 | 4 | 12 | 5,01 | 6,32 | 5,57 | 5,34 |
| CAMA | 0,23 | 136,43 | 4 | 20 | 5 | 6,78 | 5,42 | 2,42 | EMPRESA | 0,06 | 115,89 | 7 | 2 | 5,71 | 5,03 | 5 | 8,96 |
| PLAYA | 0,23 | 41,96 | 5 | 5 | 6,29 | 6,56 | 6,18 | 3,2 | PLATA | 0,07 | 47,68 | 5 | 6 | 6,16 | 6,32 | 4,04 | 6,76 |
| RUIDO | 0,14 | 57,5 | 5 | 6 | 5,34 | 5,37 | 6,05 | 4,42 | SUELO | 0,07 | 148,04 | 5 | 7 | 5,26 | 6,08 | 5,81 | 3,14 |
| COLOR | 0,14 | 128,75 | 5 | 4 | 5,92 | 5,14 | 5,68 | 2,84 | SOLDADO | 0,08 | 23,93 | 7 | 2 | 5,64 | 6,23 | 4,84 | 5,86 |
| **TOTAL** | **0,19** | **73,91** | **5,63** | **5,00** | **5,87** | **5,91** | **5,74** | **3,72** |  | **0,07** | **70,45** | **5,50** | **4,25** | **5,54** | **5,94** | **5,02** | **5,58** |

Note: BAS=backward association strength; FRE= lexical frequency; L= length; N= semantic neighbors; IMA= imaginability; FAM= Familiarity; AoA= age of acquisition; ABS= abstract; CON= concrete. HI=high association; LOW= low association.

To create the context of different BAS conditions, we manipulate this measure to build four different conditions, such as, a) Abstract - high BAS (the critical word was abstract and the words of the lists were strongly associated with it); b) Abstract low BAS (the critical word was abstract and the words of the lists were weakly associated with it; c) Concrete high BAS (The critical word was concrete and the words of the lists were strongly associated with it; d) Concrete low BAS (the critical word was concrete, and the words of the lists were weakly associated with it). The set of lists was counterbalanced using a *Latin-square* approach, which allowed each *list* to act as both a stud*y* and a distracted material.

### Procedure

Up to three participants sat together in one room in front of 17’’ CTR computers’ monitors. They were positioned so that they could not see the other participants’ screens. Headphones were provided as protection against potential noise. Stimuli were presented in black letters (font-size: 18) on a white background. Each participant studied 16-word lists and the presentation order of the lists was counterbalanced. The participants read specific instructions at the beginning of each phase. The study phase started with the word “list” and its corresponding number centered on the screen, to make sure that the participants would focus on the stimuli material. This word remained on the screen for 3000 ms. and was followed by the twelve words from the respective list. Each word was shown individually for 2000 ms. After a break of one second, the name of the next list appeared, followed by its content. This procedure was repeated until all 16 lists were presented to the participant.

Immediately after the study phase, a distractor task was presented to the participant. This task consisted in deciding whether several mathematical equations were correct or false. The distractor task was presented for 120 seconds in all the cases. The participants received feedback after each decision and the time limit for each decision was set at 100 seconds.

Finally, the participants were asked to complete a test phase, where the participants had to recognize a pool of words as studied or not studied (new). The questionnaire contained a total of 96 words formed of 16 critical words, 32 studied words (words no. 2 and 7 of each studied list), 32 distractors (words no. 2 and 7 of each non-studied list), and 16 critical control words (critical words associated with the non-studied lists). Each of the 96 trials began with a fixation cross displayed for 500 ms. Afterward, a word was presented on the center of the screen and the participant had to decide whether that word had been studied before or not by pressing the keys “A” (“No.”) or “L” (“Yes.”). The word remained visible for the participant until a response was given (i.e., it was self-paced). After the participants’ response, a new trial began. Response keys were counterbalanced across participants.

Figure 1 depicts a schematic view of the procedure.

**Figure 1.**
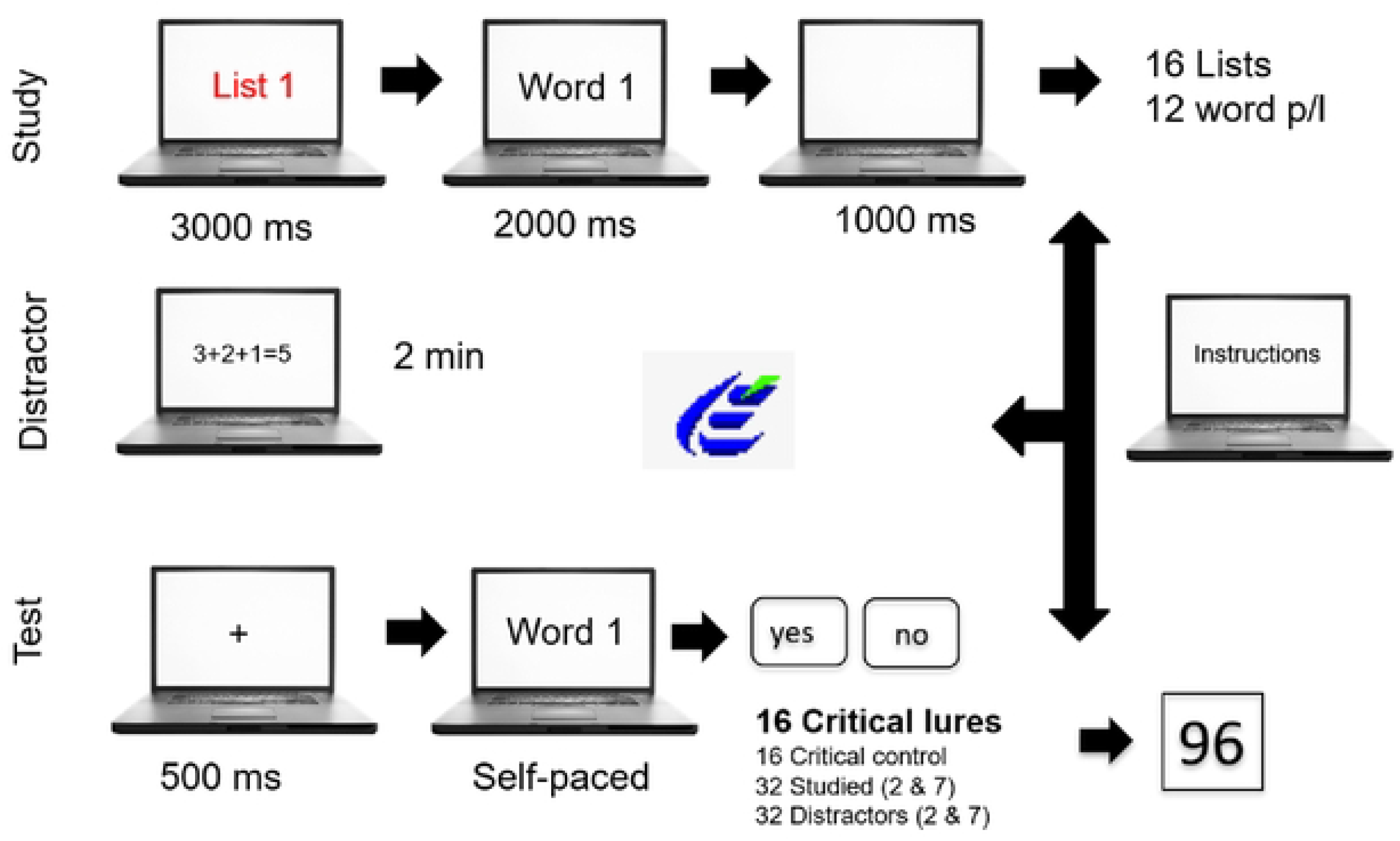
Experiment procedure.

## Results

All participants performed the distractor task correctly and no one got a score under 80%. The statistical analysis was performed only on participant’s “yes” responses in the recognition task.

### Correct Recognition

To test correct recognition, a one-way ANOVA with repeated measures was performed comparing the different types of words: critical, critical control, distractor, and correct. Results showed that type of word was significant [*F* (23) = 138.89, *p*<0.001]. Pairwise comparisons (Bonferroni) demonstrated that the proportion of “yes” responses for “studied” items (*M* = 0.75; *SD* = 0.07) was higher than for both “distractors” (*M* = 0.15; *SD* =.011) and “critical control” (*M* = 0.26; *SD* =.16) words (all *p* < .0001). No differences were found between correct and false recognition (*p* = 0.44)

### False Recognition

The mean proportion for “yes”-responses to critical lures was 0.68 (*SD* = 0.15). To test the relationship between concreteness and this false recognition effect (that was as high as the correct recognition), an ANOVA with repeated measures was performed. Results revealed a statistically significant interaction [*F* (1,23) = 4.96; *p* < 0.05] between concreteness and BAS, indicating that false recognition of critical words was higher for abstract words in the high BAS condition (*M* = 0.80; *SD* = 0.24) than for abstract critical words with low BAS (*M* = 0.63; *SD* = 0.29), as well as for weakly associated concrete words (M = 0.66; SD = 0.23) and concrete words with high BAS (M = 0.65; SD = 0.30) (see Fig. 2). No other effect reached statistical significance.

**Figure 2.**
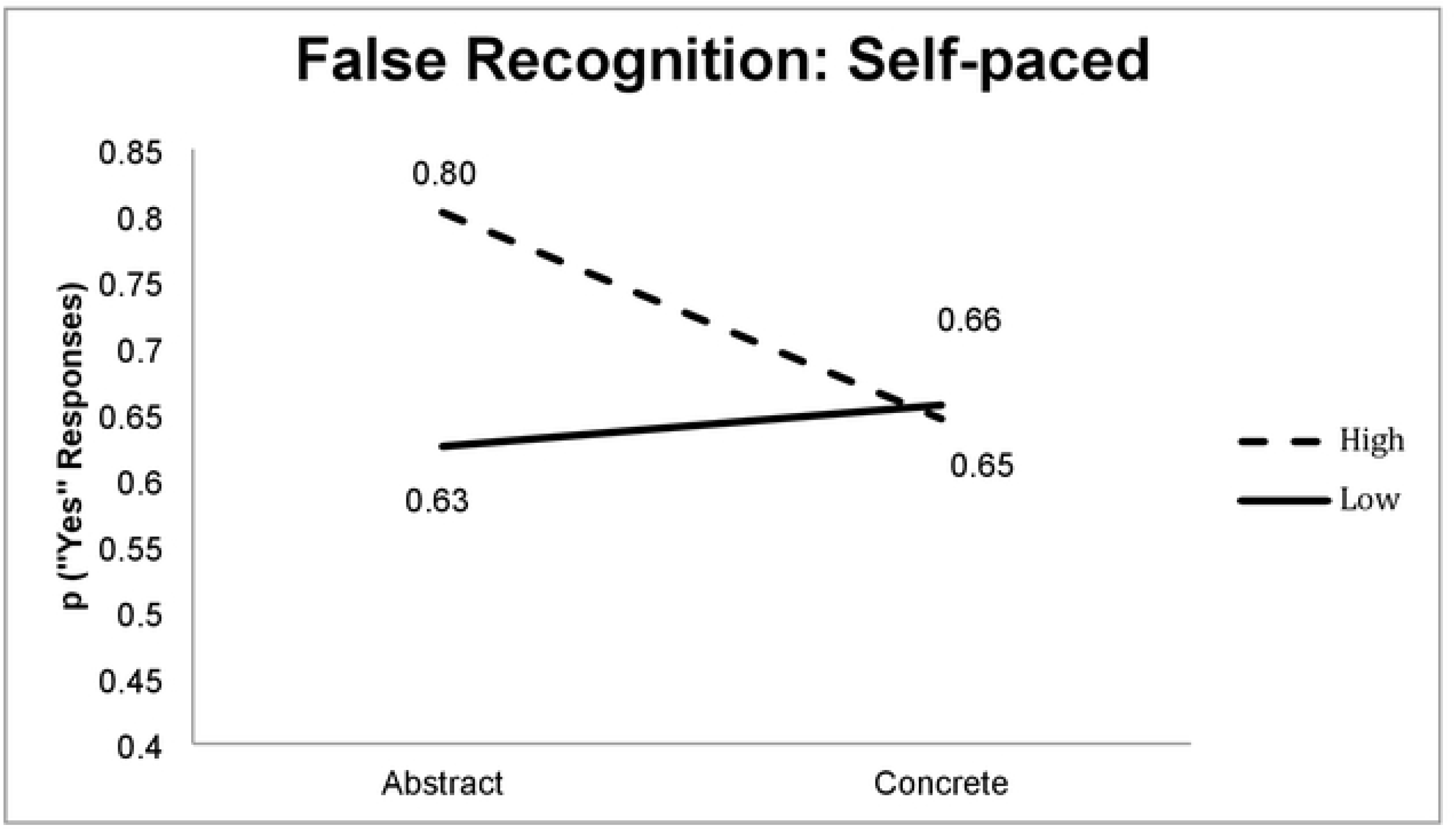
False memory effect: self-paced

## Discussion

The main goal of this experiment was to explore the impact of the relationship between word concreteness of the critical lure and the backward association strength between this word and the list of its associates over false recognition. Interestingly, we found that false recognition was higher for those abstract lures that were strongly associated with their list. These results are in concordance with Crutch et al (2006) and Duñabeitia, et. al (2009) and are in favor of QDR. From the psycholinguistic perspective, these results are also consistent with the idea that abstract and concrete concepts have a difference in processing, leading concrete words to be more accurate and faster to be processed. This is, a high number of false memories (in the case of abstract words), could be showing a less effective editing process to filter them out. Thus, the association will be the mechanism that participants use to accept an abstract not presented word as presented.

Two important considerations should be done in this part of the study. On the one hand, associative activation is a relatively short-term phenomenon that takes a few minutes before fading away. In this case, the question of the activation being present at the test phase is an important issue many authors have considered before.

A simple way to determine the role of associative activation in false recognition is by decreasing the way it acts at test minimizing the amount of time participants will have to produce the answer. This will lead to a quick study/new judgment that will be based almost entirely on processes of familiarity. In this case, not only one can expect increases in false memories but also a decrease in correct recognition. In this scenario, the question that arises is: will false memories show a concreteness effect? That was the reason for creating the second experiment.

## Experiment 2

In experiment 1, we found that association strength modified the false recognition effect of abstract words only. As mentioned before, the idea of this second experiment is to explore the effect of accelerating the response time for the participants to give a response. If participants will use associative strength to edit differently the errors for both abstract and concrete words, it should be plausible to expect a modulation in the concreteness effect. Thus, the idea of the encoding phase being a key part of false memory creation will be strengthened. In this case, the hypothesis would be that in the case of a familiarity-base judgment, associative activation will be determinant, if not, the only factor that will burst the production of false memories. Also, the increase will be affecting more abstract false recognitions since association seems to be the main organizational principle for such concepts.

## Method

### Participants

24 psychology students (19 females and 5 males participated in this study. Their mean age was 19,50 years.

### Design and materials

Again a 2 × 2 within-participants design was used with associative strength (high BAS vs. low BAS) and concreteness (abstract vs. concrete) as variables. The same materials as in Experiment 1 were used.

### Procedure

This experiment was conducted in almost the same manner as the previous one. The only change was made in the test phase. Instead of responding at their own pace, participants were instructed to respond before the target word would disappear from the screen, i.e., within a time limit of 750 ms. Feedback appeared if they did not react in time, indicating that they had to be faster. A sign to respond was provided.

## Results

### Correct Recognition

Again, an ANOVA with repeated measures was performed to check for the correct recognition effect. Results revealed a main effect of type of word [*F* (23) = 47.95, *p* < 0.001]. Pairwise comparisons showed that this effect was driven by more “yes”-responses for “studied” words (*M* = 0.49; *SD* = 0.13) and for critical lures (*M* = 0.58; *SD* = 0.13) in comparison to “critical control” words (*M* = 0.35; *SD* = 0.21) and distractors (*M* = 0.21; *SD* = 0.16).

### False Recognition

An ANOVA with repeated measures failed to show a significant main effect of concreteness or BAS and neither showed any interaction between concreteness and BAS. However, there was a tendency for abstract words in both high and low BAS to be falsely recognized in comparison to concrete words of both types. (See Fig. 3).

**Figure 3.**
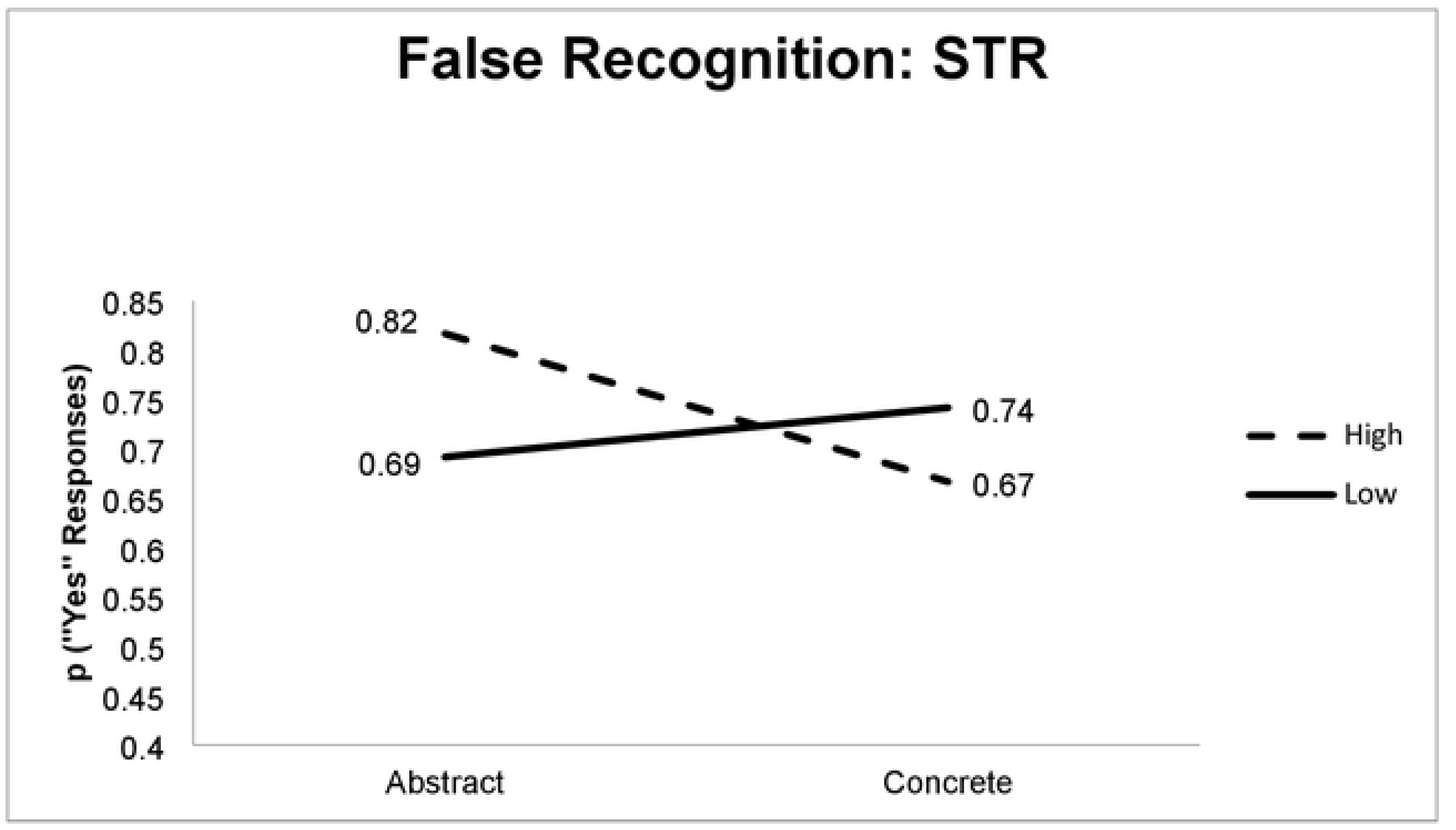
False memory effect: fast response

## Discussion

The main goal of this experiment was to explore the role of time pressure at test for creating abstract and concrete false memories. First, the first analyses provide interesting insights: false alarms and correct recognition did not differ, in fact, numerically, there was a higher proportion of critical false alarms than correct recognition. In the case of the concreteness effect at critical false recognition, statistical analyses failed at showing significant main effects nor the interaction between abstract and concrete critical lures.

The more straightforward explanation of these results is that an associative activation is a powerful tool for creating false memories, and in the absence of time to edit the response at the test, it can take over the performance to base the participant’s judgments on the familiarity process only. In this case, not only abstract but also concrete critical lures are affected and any different pattern will disappear. This is, the advantage in edition concreteness false alarms disappeared in the case familiarity-base judgments are encouraged.

Another possibility is that the concreteness effect is there, especially if we look at the numerical performance of the participants (fig. 3), but the statistical power is not enough to reveal those differences. However, this experiment is showing that noisy and less accurate performances are present when time pressure is applied and thus, participants are using associative strength to perform better at the test when recollection details are not available.

To better address this point, the next step will be to explore the neural mechanism of associative activation at the test. Thus, convergence evidence will help us to provide a more complete explanation of the concreteness effect in the context of false memories.

## Experiment 3

The objective of this experiment was to examine the neural correlates of false memories and their relationship with the behavioral results found in earlier experiments. The concreteness exploration in the brain has produced clear results about the neural mechanisms that underlay this difference. fMRI studies, for example, have shown that there seem to be neural correlates that differ from abstract and concrete concepts. Specifically, Binder et al., (2005) have found that there were areas in the left lateral temporal lobe that were equally activated by both concept types, whereas bilateral regions including the angular gyrus and the dorsal prefrontal cortex were more engaged by concrete words. In the field of the temporal domain, EEG has been used to test the time curse of abstract and concrete activation. Different authors have found that event-related brain potentials (ERPs) for concrete words usually show a long-lasting negativity (also called N700) compared to ERPs for abstract words (Barber et al.2013; Holcomb et al., 1999; Huang et al., 2010; Kanske & Kotz, 2007; Kellenbach et al., 2002; Kounios, 2007; West & Holcomb, 2000). However, there have been also results that have found an earlier wave, peaking around 400 ms. that is different for both abstract and concrete words. This N400 effect is crucial since this time window has been frequently associated with semantic processing and candidate activation (Rabovsky et al. 2018; Rabovsky et al. 2012; Rabovsky & McRae, 2014). The main goal of this experiment was to unravel the neural effect of studying word lists associated with abstract and concrete words.

## Method

### Participants

A total of 24 psychology students (8 men) took part in this experiment. The mean age was 20.50 (*SD* = 2.12) and all of them had normal or correct-to-normal vision. Participants were informed about the experiment’s characteristics and were asked to sign a consent form.

### Materials

Same as exp. 1

### Procedure

Before the start of the experiment, participants were seated in a quiet room and then they were fitted with an EEG recording cap. After that, participants were instructed to memorize sixteen twelve-word lists for an upcoming memory test. Each list was presented word by word and each word was presented for 2 seconds. Right from the beginning, the list label was presented for 5 seconds. The test phase was immediately presented after participants finished a 2-minute interference task. Each trial started with a fixation cross that remained on the screen for 1 second, followed by a 500 ms. blank screen and the memory probe, that lasted 1000 ms.

Subsequently, a blank screen was presented for 2 seconds, and later participants were asked to give a “yes/no” response to the memory probe with no time pressure imposed.

### EEG recording

Scalp voltages were collected from a 60 Ag/AgCl electrode setting, which were mounted on an elastic cap (*ElectroCap International, Eaton USA 20-10 system*) referenced to the earlobes. Four additional electrodes were used to monitor eye movements and blinking. Inter-electrode impedances were kept below 10 KΩ. EEG was filtered at 120 Hz and digitalized at a sampling rate of 500 Hz. Offline, the EEG was transformed to average reference, and later eye movements and blinks were corrected using an ocular correction method (Gratton & Coles, 1983). A 0. 032-30 Hz offline filter was applied before segmenting the EEG to obtain epochs extending from 200 ms before to 1700 ms. after the stimulus onset (baseline correction was from - 200 to 0). Artifact Rejection was performed in a semiautomatic way and a maximal voltage of 50 µV/ms. was permitted.

## Results

### Behavioral results

An ANOVA with repeated measures comparing concreteness (abstract vs. concrete) and BAS (high and low) showed that the abstract critical words generated higher rates of false recognition in comparison with the concrete ones [*F* (1,22) = 6.58; *p* <0.05]. The *F* test performed contrasted the simple multivariate effect of concretion in each BAS level and reflected that the proportion of “yes” answers given to high BAS critical words was higher compared to those given to abstract critical words of low BAS [*F* (1,22) = 7.30; *p* <0.05] (see figure 4).

**Figure 4.**
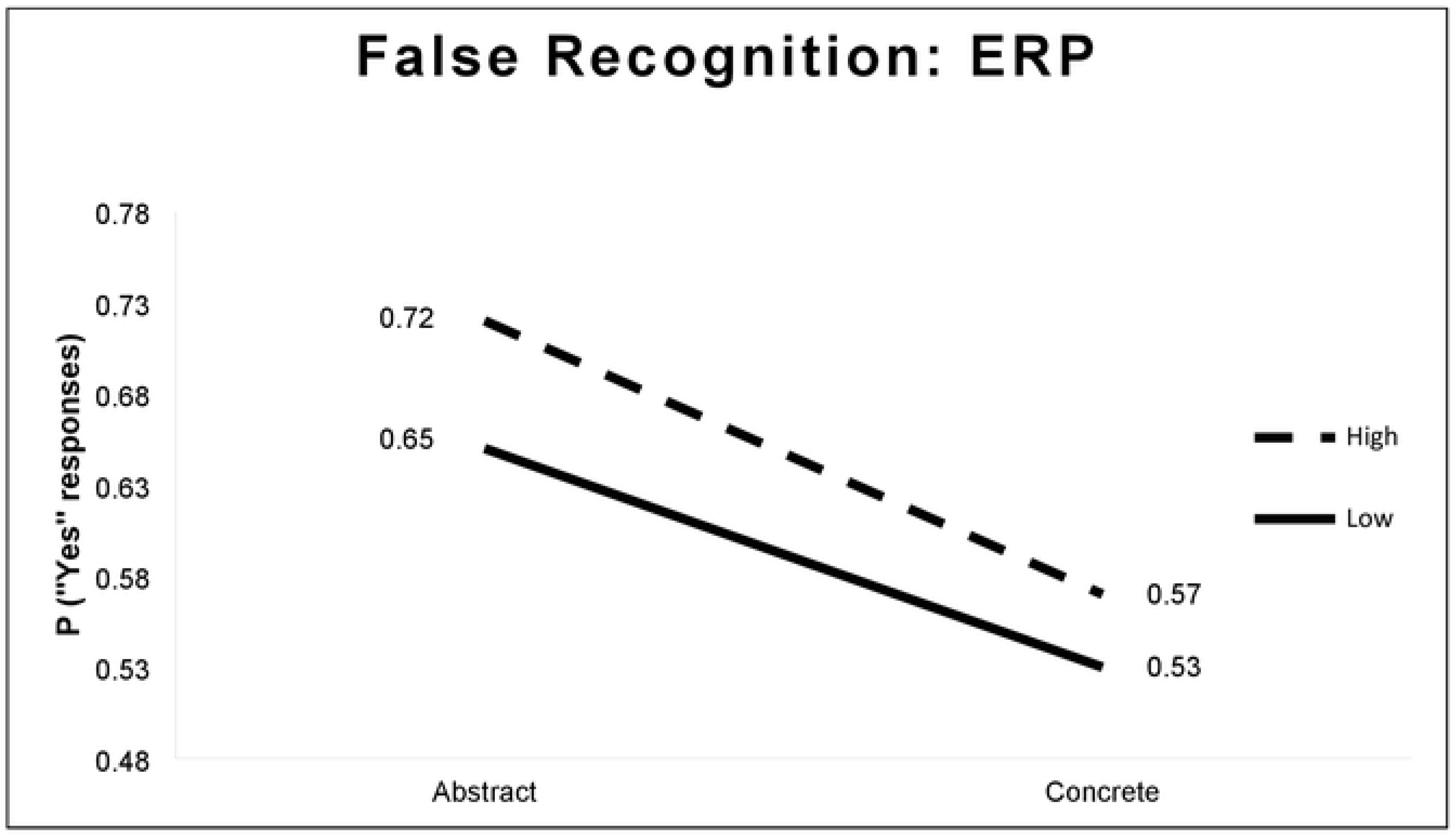
False memory effect: ERP

### Encoding phase

Segments recorded in the encoding phase were sorted out according to participants’ latter elicitation/not elicitation of the critical lure. Global field power was used to average across all the electrodes. One participant was removed from the final analysis due to the lack of valid segments in one of the 2 experimental conditions. Because different lists induced memory illusions at different rates across participants, additional segments in one condition were removed randomly, so that conditions were parallelized.

### N400

An ANOVA, with elicitation (elicit vs no-elicit), concreteness (concrete vs abstract) and BAS (high vs low) in the activity obtained in Cz electrode and in the time-window 250-350 milliseconds showed a triple interaction, statistically significant among elicitation, concreteness, and BAS [*F* (2,10) = 4.63; *p* < 0.05), reflecting those words highly associated to abstract lures showed a reduced N400 amplitude in comparison with the rest conditions but only in the elicitation condition (See Fig. 5). Since the distribution of this effect was central, the CZ, electrode was chosen to present the effect graphically.

**Figure 5.**
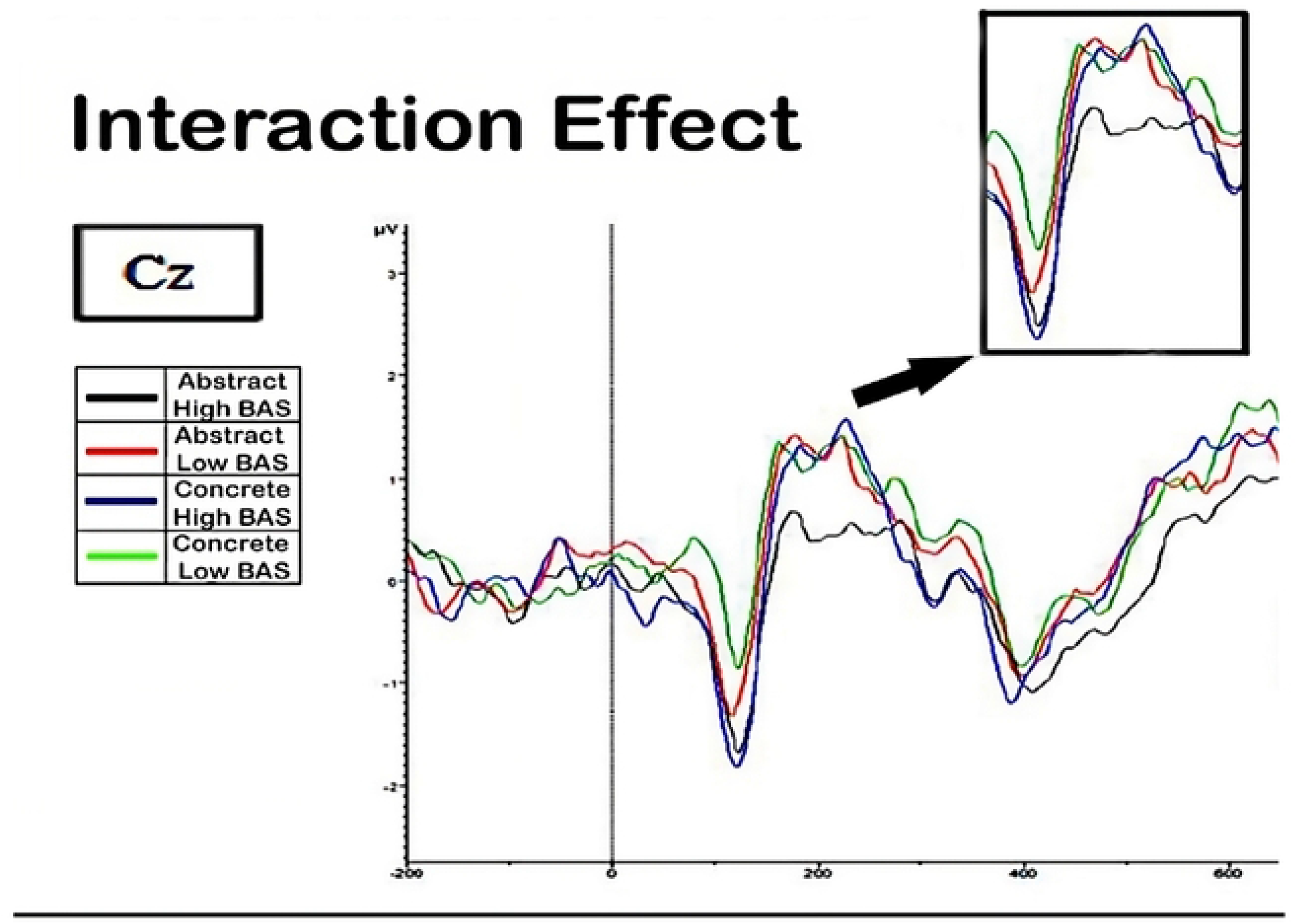
False Memory Effect: early N400

## Discussion

The main goal of this experiment was to explore the neural mechanisms of false recognition in both abstract and concrete words. Specifically, we focus on the encoding test, and we created a condition where we could compare both eliciting and not eliciting false memories. The idea behind this approach was that differences may arise when studying lists that will produce a false memory. We remark on three important issues in this experiment. First, we found an early N400 concreteness effect, which we can interpret as an effect of the differential associative activation.

Indeed, the N400 component has been considered a neural marker of semantic processing. It is often induced by words or sensations that are incongruent or semantically unexpected within a specific context. While the N400 is not directly associated with false memories, it has been researched in concreteness and semantic processing, which may indirectly relate to false memory generation.

In fact, research has demonstrated that concrete words often evoke smaller or less negative N400 amplitudes compared to abstract words. This trend shows that concrete words benefit from facilitated semantic processing compared to abstract words (Kutas, & Federmeier, 2011; West, & Holcomb, 2002). Although our data show a decreased N400 effect for the abstract high associated lists, this effect is not exactly a concreteness effect since it comes from concrete and abstract words that are associated with a critical not presented word. One way to interpret these contrary results is in line with the Associative Activation theory, which stablishes that association is an important organizational principle for abstract words. This early N400 effect can be showing that at encoding, associative activation could be used to enrich the semantic processing that will lead to a false recognition just for the abstract words.

## General Discussion

Our main goal in this study was to evaluate the role of word concreteness on false memories by using one of the most widespread tools which allows doing so in a straightforward and controlled way: the DRM paradigm. In the first place, we ran a standard experiment giving participants enough time to respond at their own pace. The results of this first experiment showed that abstract words generated more false recognition, especially if BAS is high. This result fits perfectly with the QDR theory (Crutch & Warrington, 2005) because the association is crucial for processing abstract but not for processing concrete concepts.

Regarding association, it is important to clarify that in QDR theory this term refers to the link that exists between two concepts that are either similar in meaning or tend to co-occur in language. These concepts are stored in semantic memory and depending on the nature of the link they can form a qualitatively different semantic network. Thus, based on theories of spreading activation, the processing of a concept activates not only its representation but also the representation of related ones (Collins & Loftus, 1975). Theories that explain the formation of false memories often refer to this model to explain why a non-presented word is included in the set of “yes”-responses when participants are asked to decide if a memory probe has been studied or not. Specifically, the Associative Activation Theory proposes that a lure is activated due to the processing of a list of the lure’s associates. However, there is still a problem related to this assumption: the activation phenomenon duration. Thanks to the vast research on language processing, the duration of the activation of the semantic network is dependent on various factors, like association strength, recency of the exposure, and the nature of the stimulation. Since we have different conditions of association strength, we can be sure that, at least in part, the BAS has influenced the creation of false recognition. Regarding the nature of the stimulation, we can also be confident in assuming that the differences in concreteness can also contribute to the false memory phenomenon. In consequence, the activation of the lure could occur at study, causing the formation of a new representation. If this one is strong enough to remain active until the test phase, participants will retrieve the episodic new representation, and in consequence, the participants will mistakenly decide that the item was part of the studied material. Additionally, the related lure could have been activated during the test phase thanks to the presentation of the associates that were studied previously. This could have caused the lure’s familiarity to increase and thus, the participant may have attributed this familiarity to a prior presentation of the memory probe.

Regarding the data collected in this study, it seems probable that activation occurred at the test because different manipulations of this phase affected our participants’ performances in different ways. It is also possible that activation stems from the study phase causing the participants to produce more false alarms for abstract than for concrete words. As the association is crucial for the first ones, there may have also been a summation of activation both at study and test that caused the effects and the trend we found in the three experiments in favor of abstract words. The ERP study seems to confirm this last assumption as the study phase was explored subsequently. Firstly, behavioral data was in line with our previous experiments. Secondly, when participants were studying word lists that eventually generated a false alarm, but only in the abstract condition, showed more amplitude in the N400 peak than the remained conditions. Those results can be seen as a reflection of a primary activation process for abstract words, which mainly rely on association strength.

Taken together, we propose a DIM-HA effect (high-associated abstract differential illusion memory), which is interesting, as it provides another way to explain the concreteness effect from the false memory perspective. Further research needs to be done on this topic to delve deeper into this phenomenon.

In sum, this study provides convergent evidence that contributes to advance in the knowledge of the possible mechanisms that lead to the creation of false recognition, and, at the same time, adds useful evidence for the concreteness debate. The interaction between association and concreteness both from the psycholinguistic and from the memory fields will build a stronger case for the explanation of the cognitive domains of language and memory regarding the representation of concrete and abstract concepts.

## Supporting information

S1 Fig. **Experiment procedure.**

S2 Fig. **False memory: self-Paced**

S3 Fig. **False memory: fast response**

S4 Fig. **False memory: ERP**

S5 Fig. **False Memory Effect: N400**

## References

Algarabel, S. (1996). Indices de interés psicolingüístico de 1.917 palabras castellanas Psycholinguistic indexes of 1,917 Spanish words. Cognitiva, 8(1), 43–88. https://doi.org/10.1174/021435596321235298

Barber, H. A., Otten, L. J., Kousta, S.-T., & Vigliocco, G. (2013). Concreteness in word processing: ERP and behavioral effects in a lexical decision task. Brain and Language, 125(1), 47–53. https://doi.org/10.1016/j.bandl.2013.01.005

Binder, J. R., Westbury, C. F., McKiernan, K. a, Possing, E. T., & Medler, D. a. (2005). Distinct brain systems for processing concrete and abstract concepts. Journal of Cognitive Neuroscience, 17, 905–917. https://doi.org/10.1162/0898929054021102

Bleasdale, F. A. (1987). Concreteness-Dependent Associative Priming : Separate Lexical Organization for Concrete and Abstract Words. Cognition, 13(4), 582–594. https://doi.org/10.1037/0278-7393.13.4.582

Boles, D. B. (1983). Dissociated imageability, concreteness, and familiarity in lateralized word recognition. Memory & Cognition. https://doi.org/10.3758/BF03196988

Brainerd, C. J., & Reyna, V. F. (2002). Fuzzy-trace theory and false memory. Current Directions in Psychological Science. https://doi.org/10.1111/1467-8721.00192

Brainerd, C. J., Yang, Y., Reyna, V. F., Howe, M. L., & Mills, B. A. (2008). Semantic processing in “associative” false memory. Psychonomic Bulletin & Review, 15(6), 1035–1053. https://doi.org/10.3758/PBR.15.6.1035

Breedin, S. D., Saffran, E. M., & Coslett, H. B. (1994). Reversal of the concreteness effect in a patient with semantic dementia. Cognitive Neuropsychology, 11(6), 617–660. https://doi.org/10.1080/02643299408251987

Crutch, S. J. (2006). Qualitatively Different Semantic Representations for Abstract and Concrete Words: Further Evidence from the Semantic Reading Errors of Deep Dyslexic Patients. Neurocase, 12(2), 91–97. https://doi.org/10.1080/13554790500507172

Crutch, S. J., Connell, S., & Warrington, E. K. (2009). The different representational frameworks underpinning abstract and concrete knowledge: Evidence from odd-one-out judgements. The Quarterly Journal of Experimental Psychology, 62(7), 1377–1390. https://doi.org/10.1080/17470210802483834

Crutch, S. J., & Warrington, E. K. (2005). Abstract and concrete concepts have structurally different representational frameworks. Brain, 615–627. https://doi.org/10.1093/brain/awh349

Crutch, S. J., & Warrington, E. K. (2006). Partial knowledge of abstract words in patients with cortical degenerative conditions. Neuropsychology, 20(4), 482–489. https://doi.org/10.1037/0894-4105.20.4.482

Deese, J. (1959). On the prediction of occurrence of particular verbal intrusions in immediate recall. Journal of Experimental Psychology. https://doi.org/10.1037/h0046671

Díez, E., Fernández, A., & Alonso, M. A. (2006). NIPE: Normas e índices de interés en Psicología Experimental. Retrieved from http://campus.usal.es/gimc/nipe/

Duñabeitia, J. A., Avilés, A., Afonso, O., Scheepers, C., & Carreiras, M. (2009). Qualitative differences in the representation of abstract versus concrete words: Evidence from the visual-world paradigm. Cognition, 110(2), 284–292. https://doi.org/10.1016/j.cognition.2008.11.012

Duñabeitia, J. A., Marín, A., Avilés, A., Perea, M., & Carreiras, M. (2009). Constituent priming effects: Evidence for preserved morphological processing in healthy old readers. European Journal of Cognitive Psychology, 21(2–3), 283–302. https://doi.org/10.1080/09541440802281142

Gallo, D. A. (2006). Associative Illusions of Memory. Associative Illusions of Memory: False Memory Research in DRM and Related Tasks. New York, NY: Taylor and Francis Group. https://doi.org/10.4324/9780203782934

Gallo, D. a. (2010). False memories and fantastic beliefs: 15 years of the DRM illusion. Memory & Cognition, 38(7), 833–848. https://doi.org/10.3758/MC.38.7.833

Gillund, G., & Shiffrin, R. M. (1984). A retrieval model for both recognition and recall. Psychological Review, 91(1), 1–67. https://doi.org/10.1037/0033-295X.91.1.1

Holcomb, P. J., Kounios, J., Anderson, J. E., & West, W. C. (1999). Dual-Coding, Context-Availability, and Concreteness Effects in Sentence Comprehension: An Electrophysiological Investigation. Journal of Experimental Psychology: Learning Memory and Cognition. https://doi.org/10.1037/0278-7393.25.3.721

Huang, H. W., Lee, C. L., & Federmeier, K. D. (2010). Imagine that! ERPs provide evidence for distinct hemispheric contributions to the processing of concrete and abstract concepts. NeuroImage. https://doi.org/10.1016/j.neuroimage.2009.07.031

Kanske, P., & Kotz, S. A. (2007). Concreteness in emotional words: ERP evidence from a hemifield study. Brain Research. https://doi.org/10.1016/j.brainres.2007.02.044

Kellenbach, M. L., Wijers, A. A., Hovius, M., Mulder, J., & Mulder, G. (2002). Neural differentiation of lexico-syntactic categories or semantic features? Event-related potential evidence for both. Journal of Cognitive Neuroscience. https://doi.org/10.1162/08989290260045819

Kounios, J. (2007). Insights from electrophysiology: Functional modularity of semantic memory revealed by event-related brain potentials. In Neural Basis of Semantic Memory. https://doi.org/10.1017/CBO9780511544965.004

Kousta, S.-T., Vigliocco, G., Vinson, D. P., Andrews, M., & Del Campo, E. (2011). The representation of abstract words: Why emotion matters. Journal of Experimental Psychology: General, 140(1), 14–34. https://doi.org/10.1037/a0021446

Kutas, M., & Federmeier, K. D. (2011). Thirty years and counting: Finding meaning in the N400 component of the event-related brain potential (ERP). Annual Review of Psychology, 62, 621–647. https://doi:10.1146/annurev.psych.093008.131123

Mkrtychian, N., Gnedykh, D., Blagovechtchenski, E., Tsvetova, D., Kostromina, S., & Shtyrov, Y. (2021). Contextual Acquisition of Concrete and Abstract Words: Behavioural and Electrophysiological Evidence. Brain Sciences, 11(7), 898. MDPI AG. http://dx.doi.org/10.3390/brainsci11070898

Paivio, A., & Csapo, K. (1969). Concrete image and verbal memory codes. Journal of Experimental Psychology, *80*(2, Pt.1), 279–285. https://doi.org/10.1037/h0027273

Pérez-Mata, M. N., Read, J. D., & Diges, M. (2002). Effects of divided attention and word concreteness on correct recall and false memory reports. Memory, 10(3), 161–177. https://doi.org/10.1080/09658210143000308

Rabovsky, M., Hansen, S. S., & McClelland, J. L. (2018). Modelling the N400 brain potential as change in a probabilistic representation of meaning. Nature Human Behaviour. https://doi.org/10.1038/s41562-018-0406-4

Rabovsky, M., & McRae, K. (2014). Simulating the N400 ERP component as semantic network error: Insights from a feature-based connectionist attractor model of word meaning. Cognition. https://doi.org/10.1016/j.cognition.2014.03.010

Rabovsky, M., Sommer, W., & Abdel Rahman, R. (2012). The time course of semantic richness effects in visual word recognition. Frontiers in Human Neuroscience. https://doi.org/10.3389/fnhum.2012.00011

Reyna, V. F., & Brainerd, C. J. (1995). Fuzzy-trace theory: An interim synthesis. Learning and Individual Differences. https://doi.org/10.1016/1041-6080(95)90031-4

Reyna, V. F., & Brainerd, C. J. (1998). Fuzzy-Trace Theory and False Memory: New Frontiers. Journal of Experimental Child Psychology. https://doi.org/10.1006/jecp.1998.2472

Richardson, J. T. E. (1976). Imageability and concreteness. Bulletin of the Psychonomic Society, 7(5), 429–431. https://doi.org/10.3758/BF03337237

Roediger, H. L., & McDermott, K. B. (1995). Creating false memories: Remembering words not presented in lists. *Journal of Experimental Psychology: Learning*, Memory, and Cognition, 21(4), 803–814. https://doi.org/10.1037/0278-7393.21.4.803

Roediger, H. L., Watson, J. M., McDermott, K. B., & Gallo, D. a. (2001). Factors that determine false recall: a multiple regression analysis. Psychonomic Bulletin & Review, 8(3), 385–407. Retrieved from http://www.ncbi.nlm.nih.gov/pubmed/11700893

Schwanenflugel, P. J., Akin, C., & Luh, W.-M. (1992). Context availability and the recall of abstract and concrete words. Memory & Cognition, 20(1), 96–104. https://doi.org/10.3758/BF03208259

van Schie, H. T., Wijers, A. A., Mars, R. B., Benjamins, J. S., & Stowe, L. A. (2005). Processing of visual semantic information to concrete words: temporal dynamics and neural mechanisms indicated by event-related brain potentials. Cognitive Neuropsychology, 22(3–4), 364–386. https://doi.org/10.1080/02643290442000338

Wang, J., Conder, J. A., Blitzer, D. N., & Shinkareva, S. V. (2010). Neural representation of abstract and concrete concepts: A meta-analysis of neuroimaging studies. Human Brain Mapping, 31(10), 1459–1468. https://doi.org/10.1002/hbm.20950

West, W. C., & Holcomb, P. J. (2000). Imaginal, semantic, and surface-level processing of concrete and abstract words: an electrophysiological investigation. Journal of Cognitive Neuroscience, 12(6), 1024–1037. https://doi.org/10.1162/08989290051137558

West, W. C., & Holcomb, P. J. (2002). The influence of context on semantic processing in reading. Psychophysiology, 39(4), 406–411. https://doi:10.1017/S0048577201393060

